# An *in vitro* model of breast cancer metastatic niche priming

**DOI:** 10.64898/2026.03.09.703763

**Authors:** Mia Nuckhir, Sara Cabral, Grace Eckersley, Robert B Clarke, Arti Ahluwalia, Hannah Harrison

## Abstract

Metastatic breast cancer is responsible for around 11,500 deaths a year in the UK. The primary tumour likely plays a major role in priming the distant site for metastasis and crosstalk between primary and metastatic sites may be essential for secondary tumour growth. We have developed a novel *in vitro* model in which we can further study these interactions; evaluating niche priming and cancer cell conditioning as well as assessing their influence on cell homing and colonisation.

In this paper we describe a model that we believe adds to the array of *in vitro* tools available to study various stages of the metastatic cascade, offering a unique opportunity to assess bidirectional, primary to niche interactions *in vitro*.

We show that proliferation, migration and chemotaxis, and stem cell activity are altered in both cancer cell lines and in lung epithelial cells following linked, fluidic culture. Changes in cell homing and colonisation can be modelled in cell lines and within viable lung tissue explants taken from mice, with breast cancer cells settling and growing within the lung epithelial cells and tissue explants over 7 days. The colonisation/growth of cells injected into the system closely represents that seen following tail vein injection and cancer cells can be seen to settle and grow within the lung epithelial cells.

## Introduction

Breast cancer is the most commonly diagnosed cancer in women, with over 50,000 cases per year in the UK alone^1^. Metastasis, the spread of cells from their original site to other parts of the body, typically lung, brain, bone and liver, is currently incurable. As a result, metastasis is responsible for most of the 11,500 breast cancer deaths each year^2^. The metastatic process is complex and poorly understood but is known to involve several factors, such as the ability of cancer cells to break away from the primary tumour and enter the bloodstream, their ability to survive in this alien environment whilst travelling to other parts of the body as well as to home to, invade into and colonise other tissues^3^. We urgently need to understand more about the processes by which cancers spread if we are to develop new drug targets to combat metastatic spread and/or outgrowth and improve outcome.

It has long been accepted that disseminated cancer cells must adapt to the new microenvironment at a metastatic site if they are to survive and grow and the *seed and soil hypothesis* suggests that the metastatic niche also needs to be altered to make it suitable for the cells to arrive and thrive^4^. To understand the interactions between primary and secondary site which drive successful metastasis we aimed to investigate tumour-niche crosstalk, but current *in vivo* and *in vitro* models are limited in scope and suitability for our purposes.

*In vivo* models have the potential to allow us to study the molecular mechanisms of metastasis^6^, test the efficacy of anti-metastatic drugs and to develop new strategies for preventing metastasis, but are costly, time consuming and offer very little control to assess each individual step. Cell line derived and patient derived xenografts (CDX and PDX) have proven to be a very useful offering and, in the case of PDX, have given us the opportunity to grow tumours representative of the primary tumour from which they came. PDX which are successfully grown, however, are often from specific sub-types or stages of cancer, can lose hormone receptor expression and typically metastasise infrequently^7,8^. Tail vein and intra-cardiac injections offer an alternative method to model metastasis however cell movement and colonisation is often biased towards certain tissues. Following intra-cardiac injection, cells preferentially land within the liver and lungs rather than brain or bone^9^ and, following tail vein injection, lung is the most common area of tumour growth^10^. Translation of discoveries from preclinical models to patient treatments has a high attrition rate^11^, thought to be in part due to the caveats relating to mouse use^12^.

*In vitro* models are often considered to be too basic and, although we have tight control over each phase of progression and can interrogate them in detail, the lack of different cells and relative simplicity are caveats. Recent developments in micro-fluidics have allowed the creation of more complex models in which multiple cell types can be co-cultured but remain physically separated^13^. These systems allow investigation of the interactions between cancer and normal tissue, lung cancer cells and brain cells representing a metastatic site for example^14^. Although these models have been used successfully to ask specific questions related to metastasis, there are downsides. The equipment is often manufactured in-house and is costly and the small size means a limited number of cells can be used over a short time scale. Additionally, as the cells are cultured at very close quarters, often separated only by a porous membrane, they may be more representative of already established metastases than the distant niche priming that we are focussing upon.

We have developed, and describe here, an *in vitro* model in which we can investigate interactions between the primary tumour and a distant metastatic niche. We began with a model which would allow investigation of the interactions and influence of breast cancer on the lung microenvironment, which is a site common to metastasis of triple negative breast cancer (TNBC), a sub-type which represents an area of great unmet clinical and research need^5^. Our model allows us to culture cell lines representative of the primary tumour and of the distant lung niche within individual chambers which are joined by capillaries allowing circulation of medium around each chamber. This allows signals to be passed from one cell type to another, whilst they are kept physically apart, meaning we can assess changes in both populations following linked culture. We show evidence of bidirectional signalling, with changes in proliferation rate, stem cell activity and migration occurring in both populations as well as changes with extracellular vesicle (EV) and cytokine release and see cells moving and homing from primary to metastatic niche.

To validate the system as a replacement model we have compared an *ex vivo* model of lung homing and colonisation which we use within the lab to our linked culture system. We confirm that we can mimic tail vein injection, typically performed using the PuMA model, with comparable cell growth over 7 days.

## 1. Materials and Methods

Using the *Quasi vivo™* system (Kirkstall LTD, UK), a reservoir containing 25 mL of medium is connected to a Parker Polyflex 6-Channel Peristaltic Pump (Kirkstall LTD, UK) which pushes medium around the circuit and into each of the *Quasi vivo™* QV500 chambers, each of which contains 2 mL of media (Figure 1). Chamber number is not limited but it is important to calibrate the system to calculate flow rate with each new set up (Supplemental Figure 1). In our hands, a flow of 125 µL/min offered the best rate for our cells with little shear stress occurring at the chamber base (Supplemental Figure 2, maximum 1.34 microPa, average 0.486 microPa). Proliferation, assessed by Ki67 staining (Supplemental Figure 3A), was increased at higher flow rates but remained unchanged at 125 µL/min (Supplemental Figure 3B) regardless of the chamber’s position in the circuit (Supplemental Figure 3C). The number of apoptotic cells seen was so low it was not possible to quantify but it was concluded that there was no change induced under flow (Supplemental Figure 3D). Cell morphology was also unchanged (Supplemental Figure 3E). Supplemental Figure 3A-D shows MCF7 cells but this rate was also suitable for MDA-MB-231 and 1HAEo cells in our hands. At this flow speed, the circulation time of the medium is every 3.6 hours approximately, meaning 6 full changes of medium in each 24-hour period.

**Figure 1:**
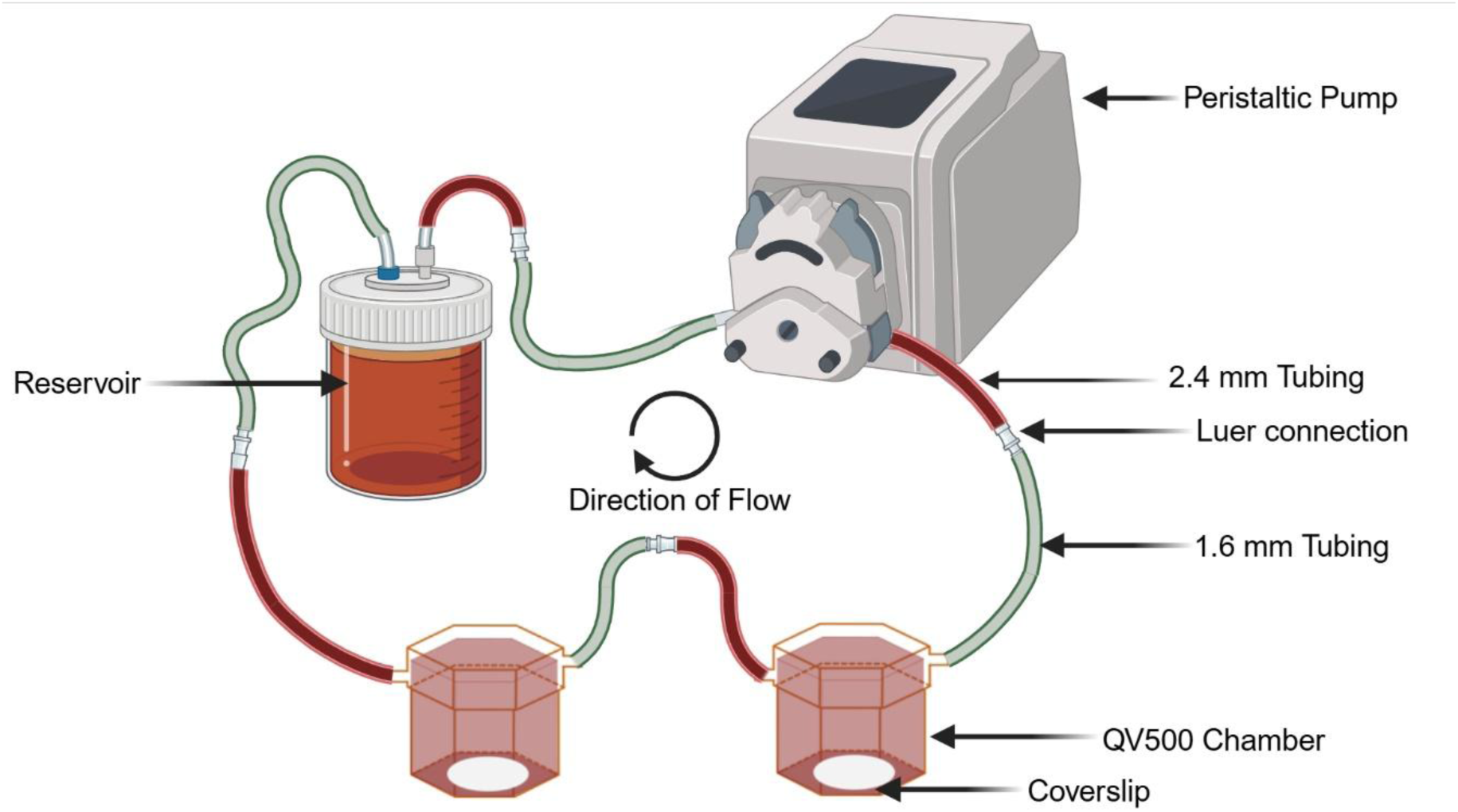
Linked culture using the *Quasi vivo*™. Medium is pumped from a reservoir and around a circuit of QV500 chambers by a peristaltic pump. Flow rate can be adjusted to achieve desired circulation speed.

Breast cancer cell lines used for this preliminary work and model development included MCF7, MDA-MB-231, both of which were purchased from the ATCC, MDA-MB-231 labelled with mApple (231-mApple), which were a kind gift from Dr Joseph Parsons, University of Manchester. 1HAEo cells, a line representing the lung airway epithelium, were a kind gift from Drs Simon Lea and Andrew Higham, University of Manchester. Cell lines were verified and confirmed as mycoplasma free. All lines were cultured in DMEM complete, consisting of DMEM High Glucose (SIGMA, D6546), 10% foetal bovine serum (FBS, Thermo, 10270106), 2 mM L-glutamine (SIGMA, G7513) and 100 µg/mL Penicillin/Streptomycin (SIGMA, P0781), and incubated at 37°C in 5% CO_2_. Sub-culture was performed when cells reached 80-90% confluence using 0.25% trypsin/EDTA (SIGMA, T4049) for 2-5 minutes at 37°C and cells were re-plated at 1:10.

NOD SCID gamma (NSG) mice were used, aged between 8-10 weeks, which were purchased from Charles River. All procedures were performed in accordance with the Animals (Scientific Procedures) Act 1986 and approved by the UK Home Office (HO Licence PP0474431) following the ARRIVE guidelines (Supplemental Table 1).

### 1.1 Immunofluorescence

Cells were plated onto coverslips (dia. 12 mm #1.5 (0.16-0.19mm), Fischer) in 24 well plates, 24 hours before transfer to the QV500 chambers to allow adherence. Cell number was optimised depending on time within the system so that the cells remained sub-confluent (e.g. for 96 hours of culture, 50,000 cells/well were plated in a 24 well plate, resulting in approximately 30,000 cells adhering to the coverslip which covered 40% of the well surface. After 4 doublings this results in approximately 480,000 cells and 95% confluence). Coverslips were then transferred to empty QV500 chambers, and the circuit was attached to the peristaltic pump before the system was primed to complete filling of the chambers and the capillaries. To assess the changes induced in breast cancer cells and lung epithelial cells following linked culture, conditioned cells were compared to naïve cells which have been exposed to the flow but no other cells.

#### Antibodies

Alexa Fluor® 488 Anti-Ki67 antibody (ab197234, Abcam) was used to assess proliferation, Alexa Fluor® 488 Anti-Prohibitin antibody (ab184813, Abcam) and Alexa Fluor® 647 Anti-Cytochrome C antibody (ab198583, Abcam) to measure viability/apoptosis, and Rhodamine Phalloidin Reagent (ab235138, Abcam) to assess cell morphology.

#### Fixing and permeabilisation

Coverslips were transferred to a 24 well plate and washed twice with PBS for 5 minutes shaking, before fixation with 4% PFA at room temperature (RT) for 10 minutes. Cells were then washed twice in PBS before permeabilisation, if required, with 0.1% Triton at RT for 15 minutes and two further PBS washes.

#### Blocking

Non-specific binding was reduced by blocking at RT for 1 hour with 500 µL 1% BSA (Merck, A2153) in PBS.

#### Staining

30 µL of primary antibody was dropped onto parafilm and coverslips flipped to bring cells into contact with the antibody for 1 hour at RT in the dark. The coverslip was then flipped back into the 24 well plate and washed twice with PBS and once with water before mounting.

#### Mounting

DAPI mounting medium (ab104139, Abcam) was dripped onto a microscope slide and coverslip flipped onto it before being left to dry overnight in the dark.

#### Imaging

Images were collected on a Zeiss Axioimager.M2 upright microscope using a 10x / 0.55 Plan Apochromat objective and captured using a Coolsnap HQ2 camera (Photometrics) through Micromanager software v1.4.23. Specific band pass filter sets for DAPI, FITC and Cy5 were used to prevent bleed through from one channel to the next. Images were then processed and analysed using Fiji ImageJ (http://imagej.net/Fiji/Downloads).

#### Scoring

Ki67 positive and negative cells were counted in 5 fields of view, containing a minimum of 200 cells, and a percentage positive score was calculated.

### 1.2 Sphere formation

To assess (cancer) stem cell activity in both the breast cancer and lung epithelial lines, non-adherent sphere culture was performed according to our published mammosphere methodology^15^. Briefly, following 96 hours in linked culture, coverslips were transferred to a 24 well plate and rinsed with PBS before trypsinisation, collection and counting. A single cell suspension was produced using a 25G needle before cells were plated at a density of 2000 cells/well in polyHEMA coated 6 well plates. 3 replicates were performed for each cell type/condition and spheres were allowed to grow for 7 days before those over 50 µm in size were counted.

### 1.3 Migration and chemotaxis

To assess changes in cell migration and chemoattraction, Boyden chamber assays were performed using cells conditioned for 96 hours within linked culture or naïve cells, exposed only to medium flow.

#### Migration

Following conditioning within linked culture, coverslips containing naïve or conditioned breast cancer cell lines were moved to a 24 well plate containing serum free medium (SFM) to serum starve for 24 hours. The cells were then trypsinised, counted and plated at cell line dependent density (MDA-MB-231 - 50,000 and MCF7 – 100,000) in 500 µL serum free medium within a Boyden chamber (Falcon Cell Culture Inserts, Pore Size 8µm, Cat. number 10136410). 700 µL complete medium was added to the well below and cells were allowed to migrate for 24 hours (Figure 2A).

**Figure 2:**
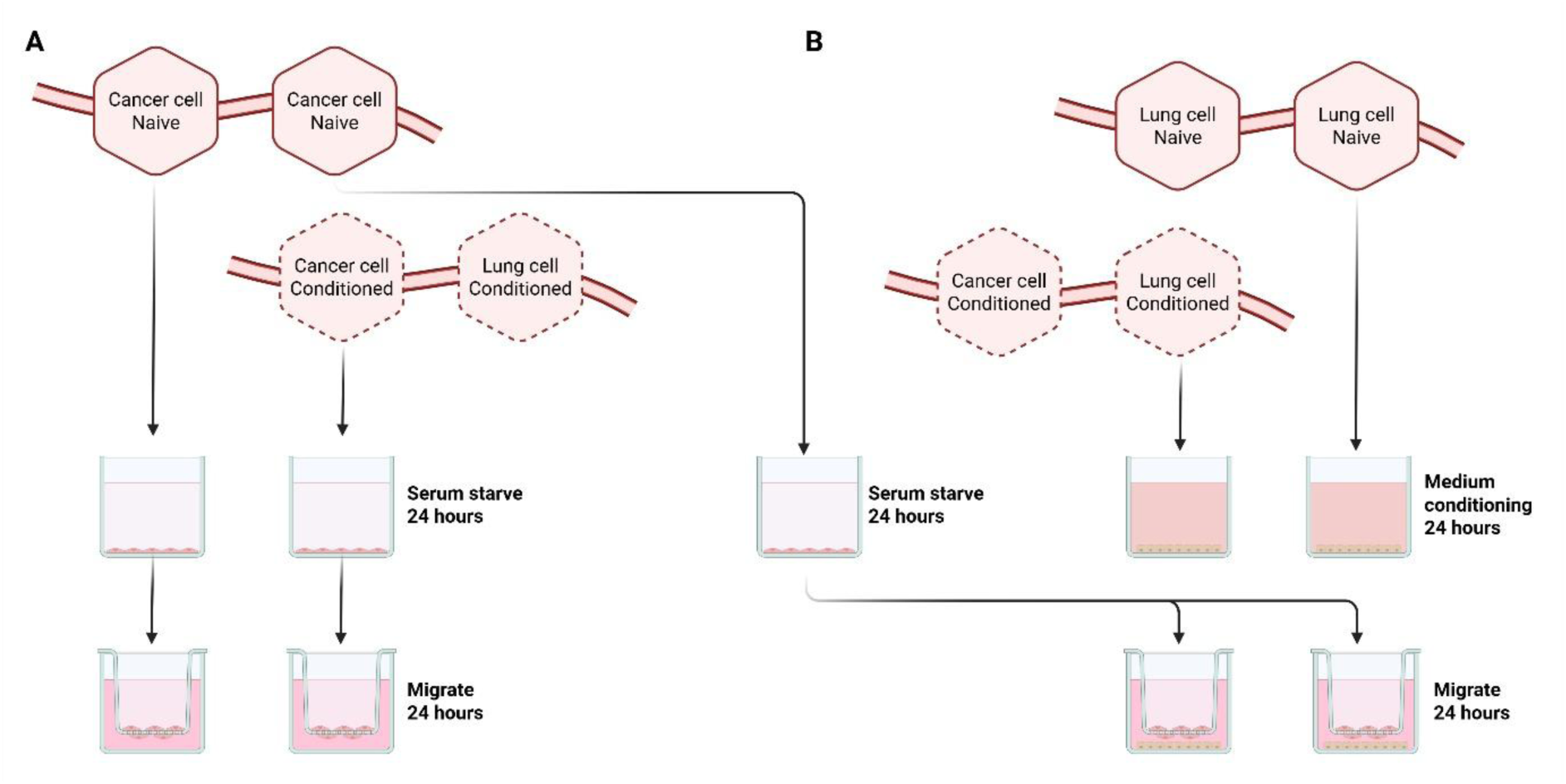
Experimental design for assessing changes in migration and chemoattraction. A) Cancer cells were grown alone (Naïve, solid lines) or in linked culture with Lung cells (Conditioned, dotted lines) for 96 hours. Naïve or conditioned breast cancer cells were then transferred into serum free medium to starve for 24 hours. The cells were then collected by trypsinisation and transferred to a Boyden chamber above complete medium and allowed to migrate for 24 hours. B) Similarly, lung cells were cultured alone (solid line) or in linked culture with cancer cell (dotted lines) before transfer to serum free culture to allow the cells to condition the medium over 24 hours. At the same time naïve cancer cells were serum starved. Naïve, starved breast cancer cells were then collected and plated into Boyden chambers above the lung conditioned medium to assess whether cancer cells were more attracted to naïve or conditioned lung cells.

#### Chemoattraction assays

Following linked culture, naïve or conditioned lung cells were transferred, on coverslips, into the well below a Boyden chamber and cultured for 24 hours in SFM. Naïve breast cancer cells, which have been serum starved for 24 hours, were then plated at cell line dependent density in 500 µL SFM within a Boyden chamber. Cells were allowed to migrate towards the lung cells, and the medium conditioned by them, for 24 hours (Figure 2B).

Membranes were rinsed in 3 changes of PBS before fixation in 4% formalin at RT for 10 minutes. Membranes were rinsed again in 3 changes of PBS before incubation in 0.1% Triton at RT for 15 minutes. Following a further 3 PBS rinses the membranes were then submerged in 0.1% crystal violet for 30 minutes at RT. Membranes were washed in 3 changes of water before drying with a cotton bud (reverse) and being left to air dry overnight. 5 fields of view were photographed and counted for each membrane to give a mean cell migration rate.

### 1.4 Signalling

#### 1.4.1 Extracellular vesicles

##### Collection from conditioned medium

For all EV work, medium was prepared using DMEM complete, containing EV depleted serum which was made in house by centrifugation of FBS at 3,000*g* for 55 minutes in a 100 kDa Amicon Ultra filter.

Following linked culture of breast and lung cells, media was collected and centrifuged at 3,000*g* for 20 minutes at 4°C to remove cellular debris. Supernatant was then concentrated in a 100 kDa Amicon Ultra centrifugal filter (Fisher, Cat. number 10781543) at 2,000*g* for 20 minutes at 4°C. The concentrated medium, approximately 200 µL, was transferred to an Eppendorf and centrifuged at 14,000*g* for 15 minutes at 4°C. Protein content was measured using a Nanodrop (Thermo).

##### Isolation and analysis of CD9, CD63 and CD81 expressing extracellular vesicles

50 µg of protein was transferred to an Eppendorf containing 50 µL of CD9 (ab239685, Abcam), CD63 (ab239686, Abcam) or CD81 (ab239687, Abcam) labelled magnetic beads and incubated overnight at RT without stirring. Next, 5 µL of the corresponding primary antibody was added (Alexa Fluor® 647 - CD9, ab187776, PE conjugated CD63, ab18236 or FITC conjugated CD81, ab239256, all Abcam) and mixed by gentle agitation. Tubes were incubated in the dark for 1 hour at 4°C without stirring. Following labelling, tubes were washed with 1 mL 2% FBS/PBS and the magnetic beads were collected using a magnetic rack (MagJET Separation Rack, MR02, Thermo). The liquid was removed from the tube carefully, so as not to disturb the beads, and then the tube was removed from the magnetic rack. Beads were resuspended in 350 µL 2% FBS/PBS for analysis on the LSR Fortessa (BD Biosciences).

#### 1.4.2 Cytokines

Following linked culture of breast and lung cells, medium was collected and centrifuged at 3,000*g* for 20 minutes at 4°C to remove cellular debris. Supernatant was then concentrated in a 3 kDa Amicon Ultra centrifugal filter (Fisher, Cat. number 11345402) at 4,700*g* for 90 minutes at RT. The concentrated medium, approximately 500 µL, was made up to a total of 1 mL and hybridised to a cytokine array membrane (RayBiotech, Human Cytokine Array C1000, Cat. Number AAH-CYT-1000) following manufacturer’s instructions. Expression was analysed using *Protein Array Analyser* plugin in Fiji Image J (http://imagej.net/Fiji/Downloads) with comparison between medium taken from flow only circuits and linked cultures.

### 1.5 Cell homing and colonisation

#### 1.5.1 Cell lines

To assess changes in cell homing, colonisation and growth, lung cells were first either conditioned in linked cultured with mApple labelled MDA-MD-231 (231-mApple) cells or remained naïve following exposure to medium flow only for 96 hours. Chambers containing conditioned or naïve 1HAEo cells were then repositioned into a new circuit and 231-mApple cells (50 cells/mL) were introduced by addition to the medium reservoir. Empty coverslips were also added to the circuit to assess random settling of cells. Cells were allowed to flow around the circuit for 24 hours.

After 24 hours, coverslips were removed, fixed and mounted (see 1.1), to assess homing, or allowed to remain within linked culture, with fresh, cell-free medium, for a further 144 hours before fixation to assess growth within the lung cell monolayers. Naïve and conditioned 1HAEo cells were also fixed before 231-mApple cells were added to assess spontaneous movement of cells during the conditioning step. 231-mApple cells were counted using “*Analyse particle*” tool in Fiji ImageJ (http://imagej.net/Fiji/Downloads).

#### 1.5.2 *Ex vivo* lung samples

##### 1.5.2.1 PuMA lung model

The *ex vivo* Pulmonary Metastasis Assay (PuMA) was established to assess the metastatic progression of human breast cancer cell lines in murine lung tissue slices^25^ (Figure 3A). On the day of the procedure, 100,000 231-mApple cells were injected into the tail vein of NSG mice. 15 minutes after injection the mice were euthanized by CO_2_ inhalation. In dorsal recumbency the sternum was removed, exposing the trachea which was then cannulated with a blunt needle. The lungs were perfused via the trachea with equal volumes of 1.2% low melting point agarose (Sigma Aldrich, A9045) and lung perfusion media consisting of 199 Earle’s Salts medium (Thermo Fisher Scientific, 31100019) supplemented with 2.0 µg/mL bovine insulin (Sigma Aldrich, 10516), 0.2 µg/ml hydrocortisone (Sigma Aldrich, H0888) and 0.2 µg/mL retinyl acetate (Sigma Aldrich, R0635) and 100 U/mL Penicillin and 0.1 mg/mL Streptomycin (Thermo Fisher Scientific, 15140122) until the lungs were fully insufflated. The lungs and trachea were removed and placed in ice cold PBS supplemented with 100 U/mL Penicillin and 0.1 mg/mL Streptomycin and incubated for 20 minutes at 4°C to solidify. Using tweezers and scalpels, transverse sections of the lung tissue were made approximately 1 mm in thickness and placed onto hemogelatin haemostatic sponges (Dentaltix, 3-271) which had been pre-incubated in a 6 well plate for 2 hours with lung culture media consisting of 199 Earle’s Salts medium supplemented with 1 μg/mL bovine insulin, 0.1 μg/mL hydrocortisone, 0.1 μg/mL retinyl acetate and 100 U/mL Penicillin and 0.1 mg/mL Streptomycin). At days 0, 3, and 7 three lung slices were removed from culture and fixed overnight in 4% neutral buffered formalin (Genta Medical, BIB10L) plus 25% sucrose (Sigma Aldrich, S7093) to maintain fluorescence. The following day the lung slices were rinsed twice in PBS and placed in a 35 mm glass bottom dish (Ibidi, 81218-200) and imaged with 5 fields of view per slice. Images were acquired on the Leica SP8 MP Upright Confocal Microscope using a 10x/0.3 Leica HCX PL fluotar objective and the 488 laser through Leica LAS X software v3.5.7.23225. Images were processed and analysed using IMARIS v9.8.0. Average mean fluorescence area was used to assess homing and colonisation as, although this is not able to provide an actual cell number, it was the most accurate and reproducible way to score the tissue.

**Figure 3:**
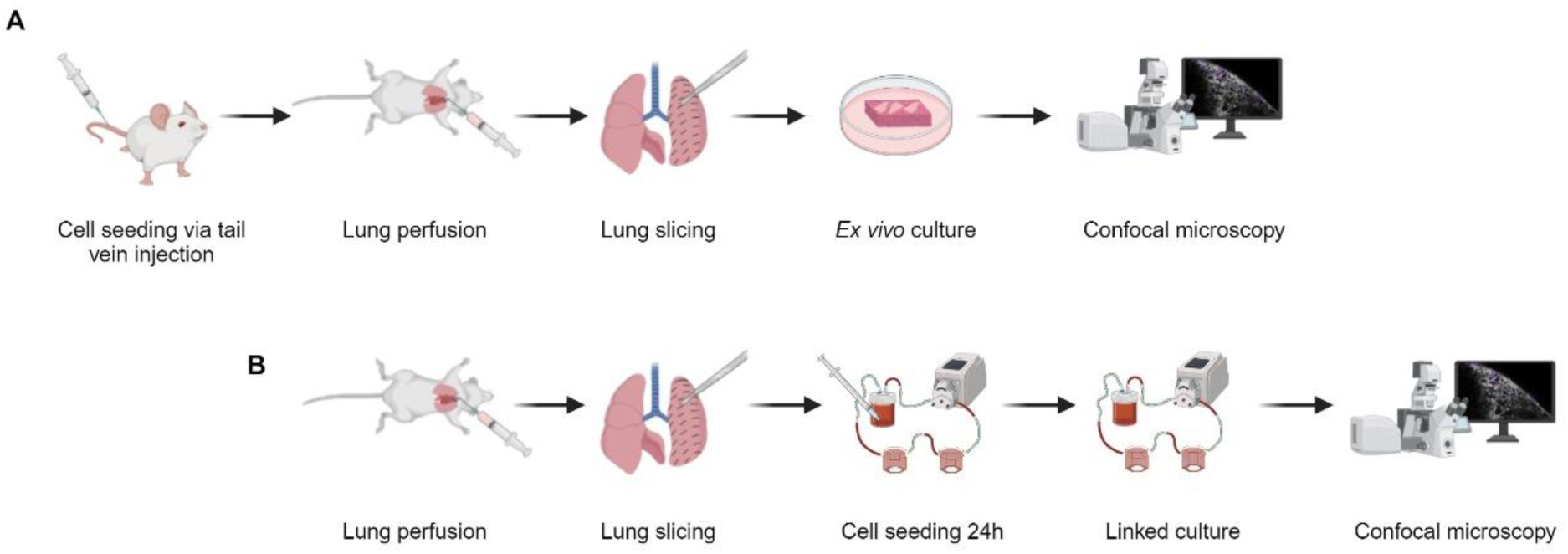
The Pulmonary metastasis assay and its adaptation to linked culture. A) 231-mApple cells are injected via the tail vein and, 15 minutes later, the lungs are perfused with agarose. The lungs are sliced and cultured ex vivo on sponges at the air/liquid interface. Cells are imaged using confocal microscopy. B) To adapt the model for use within linked culture, cell free lungs are perfused and sliced before placing within a QV500 chamber. Cells are added to the medium reservoir and allowed to circulate for 24 hours. Cell free medium is then used to culture the tissue before imaging on the confocal microscope.

##### 1.5.2.2 Homing and colonisation to PuMA slices

To adapt the PuMA model for use within our linked culture, no 231-mApple cells are injected via the tail vein but the lungs were otherwise treated as described above (Figure 3B). Slices were placed within individual QV500 chambers, above a nylon filter membrane (Fisher, 11755315) for support and below a nylon mesh with 500 µm openings (Labopolis) to hold the slices within the chamber but to allow cells to reach the tissue. For 24 hours, 231-mApple cells (5000 cells/mL) were added to the circuit, exposing the cancer cells to the lung slices. After 24 hours, the system was drained of cell containing media and replaced with lung culture media. At days 0, 3, and 7 three lung slices were removed from the system and fixed overnight in 4% neutral buffered formalin containing 25% sucrose to maintain fluorescence. Imaging was performed as described above.

### 1.6 Statistics

Where two conditions are compared, unpaired T-tests were performed otherwise, in cases where multiple comparisons were compared to control, one-way ANOVA was performed with a Dunnett’s correction for multiple comparisons. Significance was achieved at \**P*<0.05 but is also marked as \*\**P*<0.01 and \*\*\**P*<0.001.

## 2. Results

We cultured MCF7 and MDA-MB-231 cancer lines and 1HAEo lung cells in linked culture and, using the methodologies described above, assessed whether interactions between primary and secondary site occurred in linked culture. Based on our hypothesis that both sites will influence the other we predicted changes in cellular behaviour, cell hierarchy, homing and colonisation success and signalling.

### 2.1 Proliferation was altered in breast cancer and lung cell lines following linked culture

1HAEo lung cells were grown in linked culture with MCF7 or MDA-MB-231 cells for 96 hours. Figure 4A shows the rig maps for this and the following experiment (2.2). Following culture, cells were assessed for proliferation, measuring Ki67, and comparisons were made between conditioned (C) and the naïve, flow only control (N).

**Figure 4:**
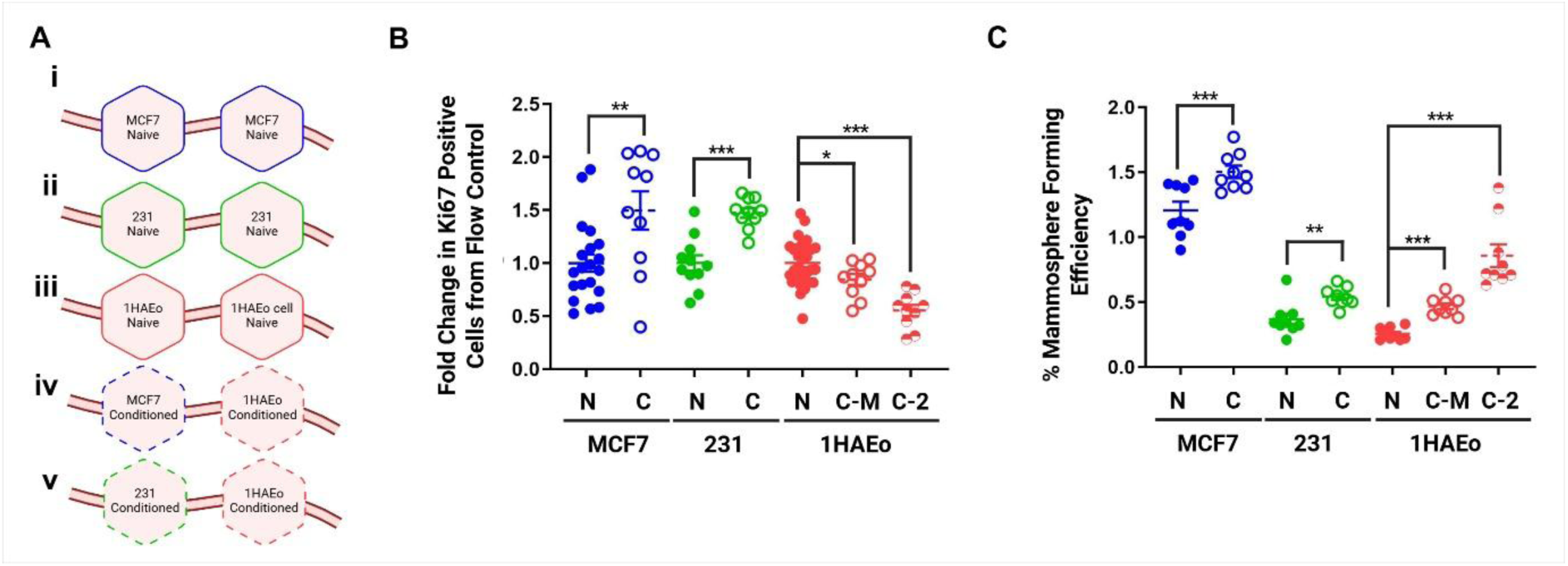
Linked culture alters proliferation and stem cell activity in breast cancer and lung cell lines. A) Rig map showing linked culture circuits used: (i-iii) Naïve MCF7, MDA-MB-231 and 1HAEo, and circuits conditioning (iv) MCF7 and (v) MDA-MB-231 with 1HAEo cells. B) Proliferation is increased in naïve (N, solid) and conditioned (C, hollow) MCF7 (blue) and MDA-MB-231 (green) but decreased in 1HAEo (red) conditioned by MCF7 (C-M) and MDA-MB-231 (C-2). C) Sphere formation is increased in all cell lines following linked culture conditioning. Unpaired T-test, \**P*<0.05, \*\**P*<0.01, \*\*\**P*<0.001.

Both MCF7 and MDA-MB-231 cells were seen to proliferate at an increased rate when cultured linked to 1HAEo cells (Figure 4B). 1HAEo cells, however, showed a significant decrease in proliferation under these conditions compared to when they are grown alone.

### 2.2 Sphere formation is altered in breast cancer and lung lines following linked culture

MCF7 and MDA-MB-231 cells were cultured alone (naïve control, N) or conditioned (C) in linked culture with 1HAEo for 96 hours before plating in non-adherent sphere culture. Spheres over 50 µm were counted after 7 days of culture. Following linked culture with 1HAEo, MCF7 and MDA-MB-231 showed an increase in sphere formation (Figure 4C). As spheres are used as an indicator of cancer stem cells (CSC) number we can conclude that this population is increasing or becoming more active in the breast cancer cells. 1HAEo cells also showed an increase in sphere forming cells following linked culture with either breast cancer cell line.

These data (2.1 and 2.2) support the hypothesis of bidirectional signalling between lung and breast cells and support the use of this platform to investigate interactions between the two sites.

### 2.3 Linked culture influences migration of breast cells and chemoattraction towards 1HAEo cells

#### 2.3.1 Migration of MCF7, but not MDA-MB-231 cells, is increased following linked culture

Following 96 hours in linked culture, conditioned and naïve MCF7 and MDA-MB-231 were serum starved for 24 hours before being allowed to migrate towards complete medium for a further 24 hours. Conditioned MCF7 (C) migrated at a significantly higher rate than their naïve controls (N) (Figure 5A). This change was not seen in MDA-MB-231 when comparing conditioned to naïve cells.

**Figure 5:**
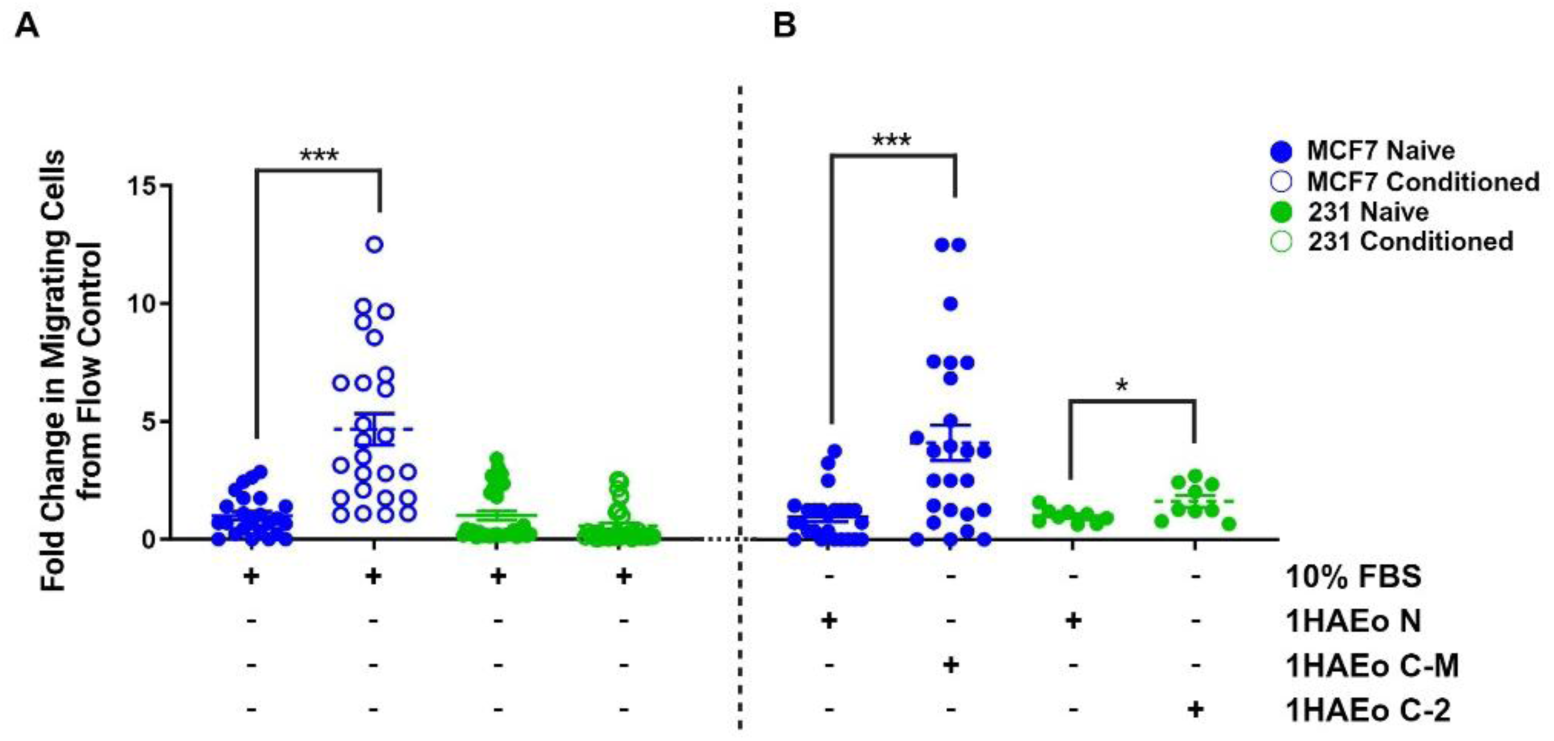
Interactions between MCF7 and 1HAEo cells in linked culture alters migration and chemoattraction. A) MCF7 cells (blue) which had been conditioned by 1HAEo cells (empty circles) migrated at a significantly higher rate than naïve cells (solid circle). No change was seen following conditioning in MDA-MB-231 cells. B) 1HAEo cells which had been conditioned with MCF7 (C-M) were more chemoattractive to MCF7 naïve cells than naïve 1HAEo cells. The same is true following MDA-MB-231 conditioning of 1HAEo (C-2). Unpaired T-test, ***P<0.001.

#### 2.3.2 Chemoattraction is increased in 1HAEo cells following linked culture with cancer cells

1HAEo cells were grown in linked culture with MCF7 or MDA-MB-231 cells for 96 hours and then used to condition serum free medium for 24 hours. Naïve MCF7 or MDA-MB-231 cells were then allowed to migrate for the 24 hours and migrating cell number was assessed. 1HAEo cells which had been link cultured to MDA-MB-231(1HAEo C-2) cells showed a small, but significant increase in chemoattraction to naïve MDA-MB-231 (1HAEo N) which had been serum starved for 24 hours (Figure 5B). A larger increase was seen following conditioning with MCF7 cells with 4-fold more MCF7 cells migrating towards conditioned 1HAEo cells (1HAEo C-M).

These findings suggest that linked culture with lung epithelial cells influences the migration and chemotaxis of cancer cells in a subtype dependent manner with luminal-like MCF7s moving more readily and having more influence upon the lung epithelium, driving it to release attractive signals into the medium.

### 2.4 Signalling

#### 2.4.1 Extracellular vesicles

Following linked culture of 1HAEo cells with either MCF7 or MDA-MB-231 cells, EVs were isolated and characterised. There was a significant decrease in CD9 expressing EVs released from linked culture of MCF7 and 1HAEo cells compared to MCF7 alone (Figure 6A). This implies the 1HAEo lung cells drive down the release of CD9 vesicles from MCF7. CD63 positive EVs are increased in linked culture of MDA-MB-231 and 1HAEo cells compared to both cell lines grown alone (Figure 6B) suggesting that production and release of this vesicle type is upregulated in linked culture. EVs expressing the surface marker CD81 were significantly increased in co-cultures of 1HAEo and MCF7 compared to either cell line alone (Figure 6C). Interestingly, in linked cultures of 1HAEo and MDA-MB-231, these vesicles decrease compared to 1HAEo alone, but increase compared to MDA-MB-231 alone showing, again, a sub-type dependent response.

**Figure 6:**
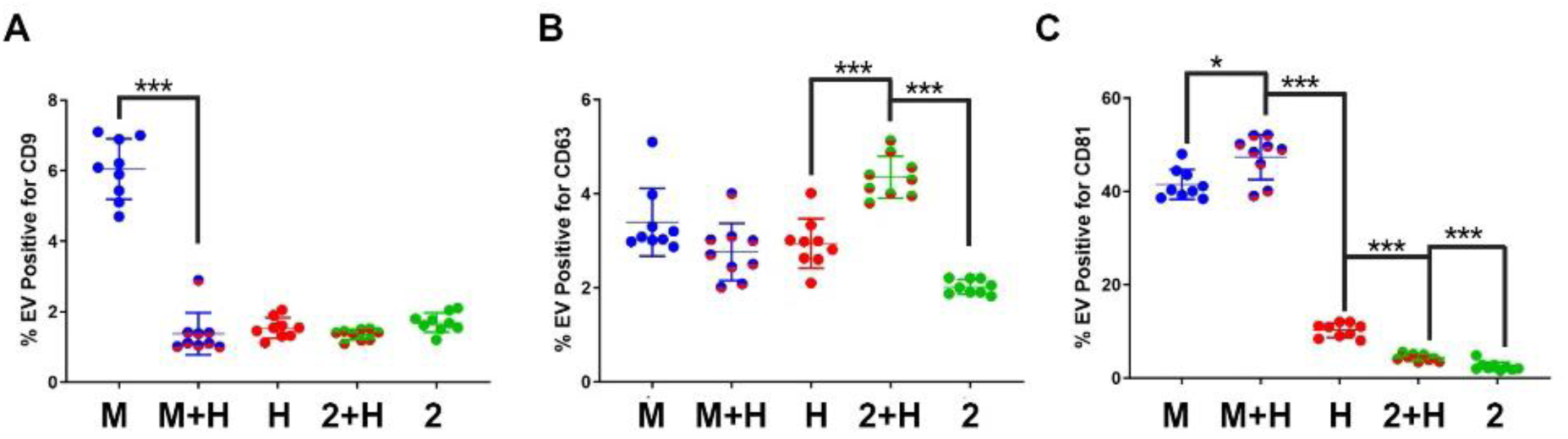
Extracellular vesicle populations change in linked culture. A) CD9 expressing EVs fall when MCF7 cells are cultured with 1HAEo cells (M-H) compared to MCF7 (M) alone and are like 1HAEo cells alone (H). B) EVs expressing CD63 are increased when MDA-MB-231 are cultured linked to 1HAEo cells (2-H) compared to both cell lines alone (2 and H respectively). C) CD81 expressing vesicles are increased when MCF7 are cultured with 1HAEo cells (M-H). In the case of MDA-MB-231 cells (2) these vesicles are increased in linked culture (2-H) but are decreased compared to 1HAEo alone (H). Unpaired T-test, \**P*<0.05, \*\*\**P*<0.001.

These data demonstrate cell line specific responses to linked culture, lend additional support for the bi-directional signalling, and demonstrate once again the importance of linked cultures. Further investigation will allow us to identify the signals within the EVs and ask what role they play in the cellular changes demonstrated throughout this paper.

#### 2.4.2 Cytokines

Whilst some inferences can be drawn from changes in EV release, such as that the increased release of CD81 EVs in each line explaining the increased stemness^16^ and increased CD9 EVs in MCF7 cells leading to increased migration^17^, further investigation is required to identify the signals within the EVs and to ascertain what role they play in the cellular changes demonstrated throughout this paper. To delve further into the signals responsible for the changes seen and to demonstrate further the usability of our model in measuring signalling between cancer and lung, we next measured cytokines within the medium following linked culture. After 96 hours in culture, conditioned medium, which had been cleared and concentrated, was hybridised to cytokine arrays and changes induced by linked culture were assessed. Cytokine expression was measured in each condition for each cell line (MCF7/MDA-MB-231 alone, 1HAEo alone and MCF7/MDA-MB-231-1HAEo linked culture) and expression in linked culture was compared to both monocultures.

In the case of MCF7 and 1HAEo cells, 53 proteins showed >1.5 fold significant increase in expression following co-culture compared to medium conditioned by either MCF7 or 1HAEo cells alone (Table 1 shows the top 30 up-regulated cytokines). Whilst in depth interrogation of these signals and their influences is beyond the scope of this method development study, most of these proteins have clear and well documented roles in migration^18–20^, chemotaxis^21,22^ and proliferation^23,24^ (highlighted as α, β and δ respectively). Others, including IL1β, have been linked to breast cancer metastasis^25^ and cancer stem cell activtiy^26^ and some, such as CX3CL1^27^, and CCL17^21^, have previously been implicated, more broadly, in metastasis. This may explain the increased migration of MCF7 cells (Figure 5A) and chemo-attraction towards 1HAEo primed with MCF7 cancer cells (Figure 5B), but this requires further investigation and confirmation. A number of cytokines which have been linked to lung metastasis were also elevated including TNFα^28^ and SDF1^29^, which play roles in lung tropism and IL1α, which is secreted by cancer cells upon lung colonisation for which it is essential^30^.

> In the case of MDA-MB-231 and 1HAEo linked culture the expression of far fewer proteins was seen - only 3 proteins showed significantly increased expression >1.5 fold. This makes pathway analysis impossible and drawing conclusions more difficult but BMP6^31^, which is upregulated, has been implicated in lung disease and in breast cancer stem cell activity so this may be an avenue warranting further investigation.

**Table 1:** Most changed cytokines following linked culture between MCF7 and 1HAEo (left) or MDA-MB-231 and 1HAEo (right). Cytokines significantly changed during linked culture are listed and highlighted as those known to relate to migration^α^, chemotaxis^β^ and/or proliferation^δ^.

| MCF7 and 1HAEo |  |  |
| --- | --- | --- |
| Cytokine | Average | P |
| Flt-3 Ligand | 2.81 | 0.0043 |
| TNF-alpha<sup>$\alpha\beta\delta$</sup> | 2.74 | 0.004 |
| IL-16<sup>$\alpha\beta$</sup> | 2.69 | 0.0045 |
| TNF-beta | 2.63 | 0.0058 |
| G-CSF | 2.59 | 0.0115 |
| CCL17<sup>$\alpha\beta$</sup> | 2.58 | 0.003 |
| IL-15 | 2.55 | 0.0062 |
| GCP-2<sup>$\alpha\beta$</sup> | 2.50 | 0.0097 |
| TGF-beta<sup>$\alpha\delta$</sup> | 2.49 | 0.0066 |
| IL-1beta<sup>$\alpha\beta$</sup> | 2.48 | 0.0109 |
| M-CSF | 2.47 | 0.0097 |
| CX3CL1<sup>$\alpha\beta$</sup> | 2.45 | 0.0134 |
| MIG<sup>$\alpha\delta$</sup> | 2.43 | 0.0097 |
| Angiogenin | 2.42 | 0.0179 |
| IL-1ra | 2.41 | 0.0126 |
| SCF<sup>α</sup> | 2.40 | 0.0068 |
| MCP-1 | 2.36 | 0.0185 |
| MDC | 2.34 | 0.0136 |
| EGF<sup>δ</sup> | 2.32 | 0.0163 |
| SDF-1<sup>$\alpha\beta\delta$</sup> | 2.31 | 0.0135 |
| BLC | 2.29 | 0.0129 |
| MCP-2<sup>$\alpha\beta$</sup> | 2.29 | 0.0148 |
| BDNF | 2.28 | 0.0143 |
| GDNF<sup>α</sup> | 2.27 | 0.0174 |
| Eotaxin-2<sup>$\alpha\beta\delta$</sup> | 2.26 | 0.0193 |
| MCP-4<sup>α</sup> | 2.25 | 0.0099 |
| FGF-7<sup>δ</sup> | 2.25 | 0.0189 |
| uPAR | 2.22 | 0.0302 |
| IL-2 | 2.20 | 0.024 |
| IL-3 | 2.19 | 0.0181 |

**Table 1: Most changed cytokines following linked culture between MCF7 and 1HAEo (left) or MDA-MB-231 and 1HAEo (right).**
| 231 and 1HAEo |  |  |
| --- | --- | --- |
| Cytokine | Average | P |
| BMP-6 | 1.59 | 0.0059 |
| IL-5 | 1.54 | 0.0183 |
| NAP-2 | 1.50 | 0.0053 |

These data importantly show that different breast cancer cell types induce differential responses from the lung cells and that the lung cells also elicit a subtype dependent response in the cancer cells to which they are linked. Also, importantly, this data demonstrate that the levels of cytokines produced from relatively low cell numbers (<500,000) can be measured in 25 mL of medium supporting the use of our model to study tumour – niche interactions.

### 2.5 Homing and colonisation

#### 2.5.1 Cell lines

Figure 7A shows the rigs and set up for this experiment. Following production of naïve (i) and conditioned (ii) 1HAEo cells, coverslips were fixed (blue) or moved to new circuits (yellow and purple boxes) with 231-mApple in the reservoir (green box). Following 24 hours of flow, cells were fixed (iii, yellow) or allowed to grow for 144 hours (iv, purple) with fresh medium containing no cells. 231-mApple were detected following pre-conditioning when no cells were injected into the medium, suggesting spontaneous movement and homing to lung cells (Figure 7B, blue boxes) but further testing is required to confirm this. It is important to note that very few cells were seen to land on empty coverslips if they were placed within the circuit suggesting that settling is selective. Cells were also visible after 231-mApple were added to the medium (yellow and purple). 231-mApple cells were seen to home to both conditions at a similar rate (Figure 7C) but over the following 6 days only those which had been previously exposed to 231-mApple cells were seen to grow.

**Figure 7:**
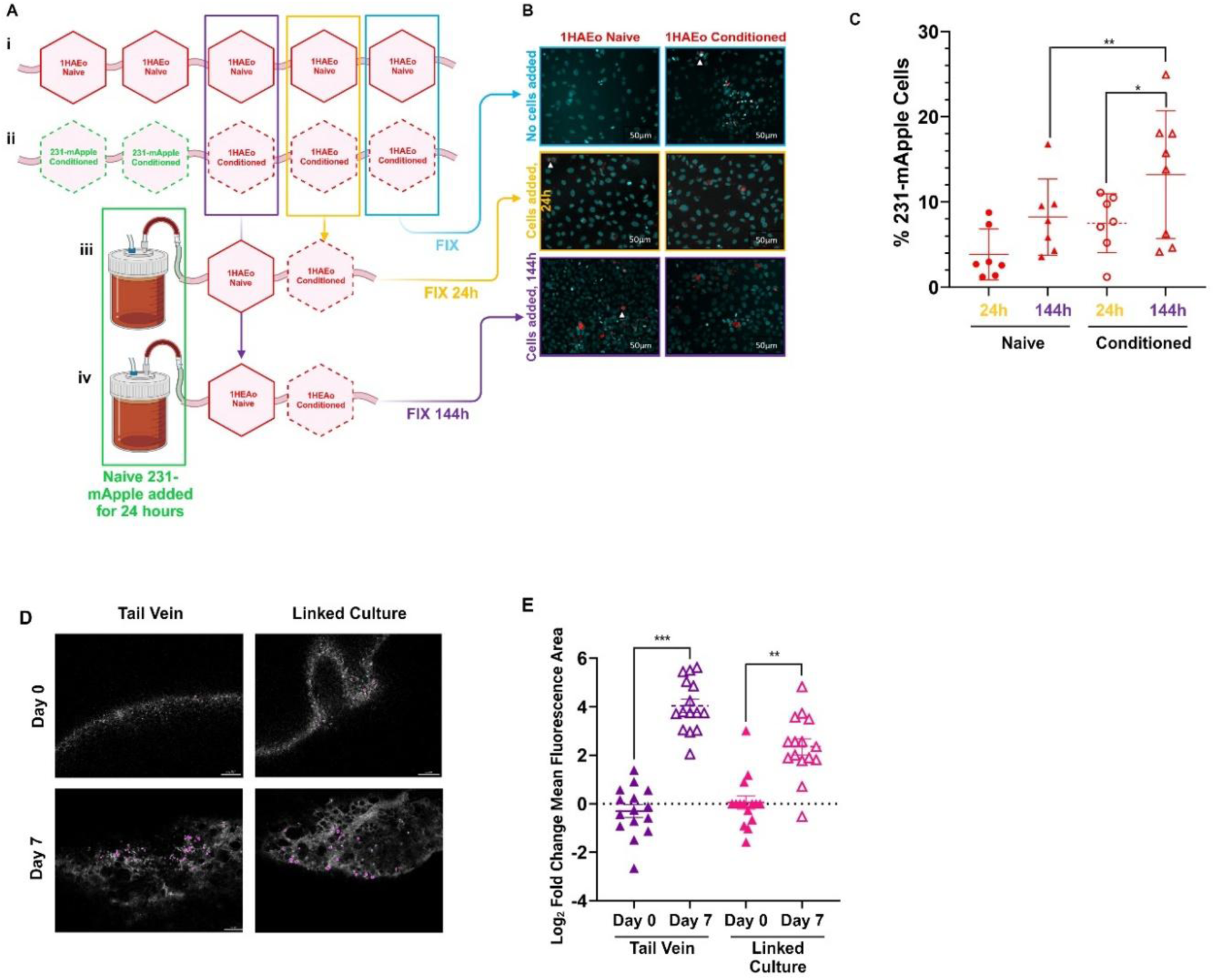
Homing and colonisation. A) Circuits used in homing and colonisation tests. i) Naïve 1HAEo cells are exposed to flow only and ii) Conditioned 1HAEo are produced by exposing the cells to 231-mApple cells for 24 hours. Cells are fixed (blue) or transferred into new circuits with 231-mApple cells in the medium (yellow and purple). iii) Cells are fixed after 24 hours or iv) allowed to grow for a further 120 hours with no cells in the medium. B) Representative images of coverslips taken from each circuit, arrows highlight 231-mApple cells (red) within 1HAEo line (blue) within naïve and conditioned 1HAEo cells. C) Percentage 231-mApple cells per coverslip was calculated. No difference is seen in settling of cells on naïve (solid shapes) or conditioned (empty shapes) 1HAEo cells, but significantly more growth is seen by 144 hours. D) Representative images taken by confocal microscopy of tissue explants. Cells are pseudo coloured purple for ease of identification. E) Significant increases are seen in both tail vein injected and linked culture added cases and these increases are comparable between methods. Log2 fold change of mean fluorescence area is shown. Scale bars represent 50 µm. Unpaired T-test, \*\**P*<0.001, \*\**P*<0.01, \*\*\**P*<0.001.

This suggests pre-conditioning of the lung niche allows cancer cells to proliferate within the niche more quickly or that without pre-conditioning the cells remain dormant within the lung. Further investigation is required to assess the cell cycle state of these cells.

#### 2.5.2 PuMA model

Using the PuMA model 231-mApple cells, which were injected into the tail vein shortly before lung collection, can be seen to settle in lung tissue using confocal microscopy (Figure 7D). Fluorescence, which is used as a surrogate for cell number, is seen to increase significantly over 7 days (Figure 7E). When tissue explants are added to linked culture and cells are added to the reservoir, cells are seen to grow over time in a comparable way with significantly higher fluorescence at day 7 (7D and E).

The findings presented here suggest that our linked culture system is suitable for assays investigating niche priming, cancer cell conditioning, homing and colonisation and that it offers a replacement/reduction opportunity for tail vein to lung metastasis experiments.

## Discussion

The data presented in this paper show the importance of more complex cultures, which take different cells/tissues into account, in metastasis research. Both breast cancer cell lines showed increased proliferation when cultured linked to lung alveolar cells whilst we see a decrease in lung epithelial cells following culture with cancer lines.

Similarly, changes in (cancer) stem cells are seen in all three lines tested, with a significant increase in non-adherent sphere formation in all cell lines. To account for the increased proliferation of the cancer cells, we allowed the naïve cancer sphere cultures to grow for an additional 48 hours in case spheres were seen to have increased in size enough to be counted (>50µm). The numbers of spheres did not increase significantly in this time (data not shown), and the colonies began to clump making counting impossible at this later time point. The lung cells, which were proliferating more slowly following linked culture, had a higher sphere count supporting our conclusion that change in sphere number equates to change in (cancer) stem cell activity/number rather than proliferation.

Migration and chemoattraction assays also showed changes in both populations following linked culture, although only by a small amount in MDA-MB-231 cells. Lung cells which had been pre-conditioned by cancer cells were more attractive to cancer cell lines than those which were naïve. This suggests that, in response to signals from the cancer cells the lung epithelial cells begin sending out signals to drive homing of the cancer cells. These signals are less influential in MDA-MB-231 than in the MCF7 cells. The migration capacity of MCF7 cells, but not MDA-MB-231, was also increased following linked culture with lung cells. This shows the importance of comparing multiple cells lines from different sub-types with different *in vitro* behaviours and different tropisms in patients. MDA-MB-231 are a highly migratory/metastatic line and this may explain the non-significant effect seen; the cells may be too migratory to influence in their normal state. Also, the line represents triple negative breast cancer (TNBC) which preferentially home to the lungs in patients. We could postulate that the signals are offering new information to the MCF7, ER-positive cells which more readily home to the bones in patients, meaning they have more influence on their behaviour.

Although we have yet to elucidate the signals involved in these behavioural changes, we can see that the release of extracellular vesicles (EV) is altered following linked culture. EVs were identified by membrane markers, CD9, CD61 and CD83. Although these markers are also known to be expressed on microparticles and apoptotic bodies^32^, the ultrafiltration of our medium, using a size exclusion column, means these smaller bodies are not collected and assessed^33^. As these vesicles, which are known to carry signals over long distances, are altered we could hypothesise that the signals being released by cancer and lung cells are changing. Although there is much to learn about the content of EVs they are suggested to have various roles in cancer; CD9 labelled EVs have been shown to influence invasion, migration and endocytosis^34^, CD63 EVs to contain Programmed death-ligand 1 (PDL1)^35^ whilst CD81 EVs have been shown to influence both invasion and tumour cell cluster formation^36^. Clearly more work is required to determine the influence of the changes that we report here.

Using our linked culture system, we show that pre-conditioning of lung cells by MDA-MB-231 makes the lung “microenvironment” more suitable for cancer cell colonisation and outgrowth. Cancer cells grow at a significantly higher rate in lung cells which are conditioned compared to naïve. Interestingly the homing and settling of cancer cells was not changed between the naïve and conditioned lung cells. This correlates with our earlier data showing that the chemoattractiveness of conditioned lung cells was not greatly increased compared to naïve cells. It does suggest, however, that there are changes induced in the niche by the primary tumour which makes it more hospitable to cancer cells which arrive there. We did not assess whether the cells which landed in naïve monolayers were in a dormant but viable state, and this will be of great importance moving forward as outgrowth is only part of the story.

Importantly our model also shows that cancer cells can spontaneously move, by lifting from their coverslip and flowing around the system. Cells did not do this to land on empty coverslips only to lung cell monolayers. Using this system will offer the opportunity to look at preference for tissue site/niche and for inhibiting cell homing.

Using our model in a direct comparison with an *ex vivo* model allows us to validate one element of its 3Rs potential which was a driving force behind its development. The pulmonary metastasis model (PuMA) has proven to be very useful in our lab to assess the colonisation of breast cancer cells within the lung following teil vein injection. We demonstrate here that we can replace the tail vein injection with addition of cells to the explant within linked culture. This offers a refinement to our PuMA model as we remove the need for the handling and injection and move straight to CO_2_ inhalation. We can culture multiple tissue explants from a single mouse, allowing us to assess the influence of several drugs or inhibitors in the post-metastasis setting and our data, not included in this manuscript, shows that we can influence *ex vivo* cell growth using neutralising antibodies to proteins expressed within the lung microenvironment. To assess the influence of drugs/inhibitors applied to cells before colonisation using the tail vein method, requires 5 mice, for example, to reach power. Using the linked culture system, a single mouse can be used, as it provides enough slices to power the experiment, offering an excellent reduction opportunity.

The model, described here and depicted in Figure 8, offers a great deal of opportunities for the field of metastasis research; it is highly adaptable, affordable, and easy to use. Our future aims are to develop the model further, incorporating different tissue explants, such as the bone, liver, and brain, as well as PDX and patient derived cancer explants. The model will allow us to investigate niche priming and its role in tropism and cancer cell dormancy as well as offering a platform on which to test drugs and inhibitors, allowing us to look in detail at their effects on different stages of the process.

**Figure 8:**
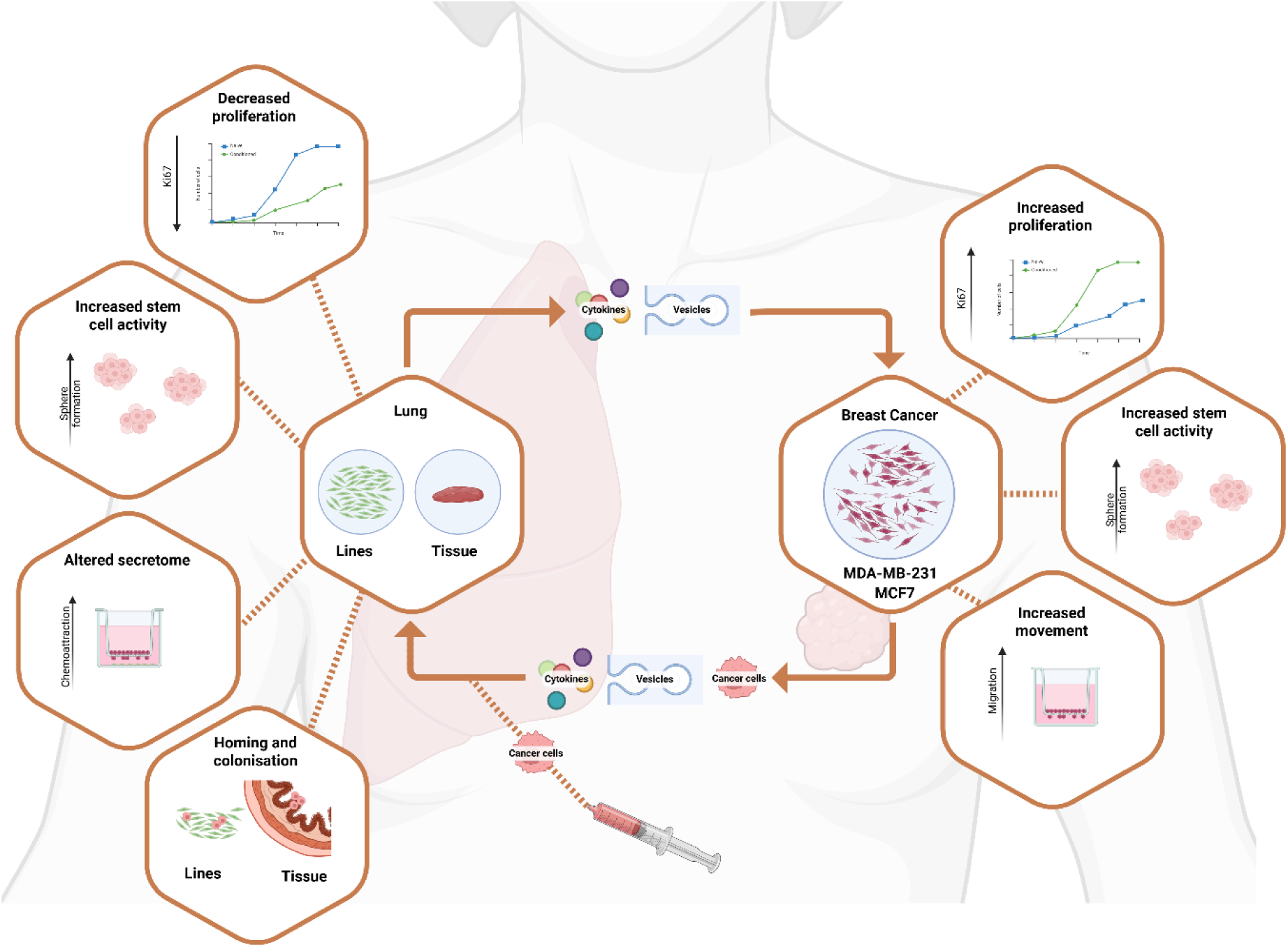
Linked culture model of niche priming. Graphical summary of the model uses in the study of breast priming and niche conditioning.

## Abbreviations

C: conditioned (in linked culture)
CDX: cell line derived xenograft
CM: conditioned medium
CSC: cancer stem cell
EV: extracellular vesicle
N: naïve (or flow only control)
PDX: patient derived xenograft
PuMA: Pulmonary Metastasis Assay
RT: room temperature
SFM: serum free medium
TNBC: triple negative breast cancer

## Author Contribution

MN - in vivo and correlation experiments, SC - linked culture experimentation and editing, GE - linked culture experimentation, RBC - intellectual input and editing, AA - intellectual input and editing, HH - conception, linked culture experimentation, manuscript writing.

## Acknowledgements

This work was funded by a joint award from the NC3Rs and CRUK, grant number NC/W000792/1 (HH), an Innovation grant from FRAME, grant number FIGUoM22 (HH) and the Medical Research Council, grant number MR/N013751/1 (MN).

The Bioimaging Facility microscopes used in this study were purchased with grants from BBSRC, Wellcome trust and the University of Manchester Strategic Fund. Special thanks go to Roger Meadows for help with the microscopy. Thanks to the Visualisation, Irradiation and Analysis team at Manchester Cancer Research Centre, specifically Steve Bagley for optimisation of PuMA image acquisition and Henry Banks for analysis. The Flow Cytometry core facility at the Manchester Cancer Research Centre was used for the EV assay and special thanks go to Antonia Banyard for guidance and input during this section of work. All figures were produced in Biorender.

## References

1 CRUK. Breast cancer statistics, <https://www.cancerresearchuk.org/health-professional/cancer-statistics/statistics-by-cancer-type/breast-cancer#:∼:text=Breast%20cancer%20risk&text=1%20in%207%20UK%20females,caused%20by%20post%2Dmenopausal%20hormones.> (2022).

2 Liu, Z. J., Semenza, G. L. & Zhang, H. F. Hypoxia-inducible factor 1 and breast cancer metastasis. Journal of Zhejiang University. Science. B 16, 32–43 (2015). 10.1631/jzus.B1400221

3 Valastyan, S. & Weinberg, R. A. Tumor metastasis: molecular insights and evolving paradigms. Cell 147, 275–292 (2011). 10.1016/j.cell.2011.09.024

4 Psaila, B. & Lyden, D. The metastatic niche: adapting the foreign soil. Nature reviews. Cancer 9, 285–293 (2009). 10.1038/nrc2621

5 Li, C. H., Karantza, V., Aktan, G. & Lala, M. Current treatment landscape for patients with locally recurrent inoperable or metastatic triple-negative breast cancer: a systematic literature review. Breast cancer research : BCR 21, 143 (2019). 10.1186/s13058-019-1210-4

6 Richmond, A. & Su, Y. Mouse xenograft models vs GEM models for human cancer therapeutics. Dis Model Mech 1, 78–82 (2008). 10.1242/dmm.000976

7 Eyre, R. et al. Patient-derived Mammosphere and Xenograft Tumour Initiation Correlates with Progression to Metastasis. J Mammary Gland Biol Neoplasia 21, 99–109 (2016). 10.1007/s10911-016-9361-8

8 Liu, Y. et al. Patient-derived xenograft models in cancer therapy: technologies and applications. Signal Transduct Target Ther 8, 160 (2023). 10.1038/s41392-023-01419-2

9 Kuchimaru, T. et al. A reliable murine model of bone metastasis by injecting cancer cells through caudal arteries. Nature communications 9, 2981 (2018). 10.1038/s41467-018-05366-3

10 Rashid, O. M. et al. Is tail vein injection a relevant breast cancer lung metastasis model? J Thorac Dis 5, 385–392 (2013). 10.3978/j.issn.2072-1439.2013.06.17

11 Fernandez-Moure, J. S. Lost in Translation: The Gap in Scientific Advancements and Clinical Application. Frontiers in bioengineering and biotechnology 4, 43 (2016). 10.3389/fbioe.2016.00043

12 Hutchinson, L. & Kirk, R. High drug attrition rates--where are we going wrong? Nat Rev Clin Oncol 8, 189–190 (2011). 10.1038/nrclinonc.2011.34

13 Xu, Z. et al. Design and Construction of a Multi-Organ Microfluidic Chip Mimicking the in vivo Microenvironment of Lung Cancer Metastasis. ACS Appl Mater Interfaces 8, 25840–25847 (2016). 10.1021/acsami.6b08746

14 Liu, W. et al. AKR1B10 (Aldo-keto reductase family 1 B10) promotes brain metastasis of lung cancer cells in a multi-organ microfluidic chip model. Acta Biomater 91, 195–208 (2019). 10.1016/j.actbio.2019.04.053

15 Shaw, F. L. et al. A detailed mammosphere assay protocol for the quantification of breast stem cell activity. J Mammary Gland Biol Neoplasia 17, 111–117 (2012). 10.1007/s10911-012-9255-3

16 Pederson, P. J., Liang, H., Filonov, D. & Mooberry, S. L. Eribulin and Paclitaxel Differentially Alter Extracellular Vesicles and Their Cargo From Triple-Negative Breast Cancer Cells. Cancers 13, 2783 (2021). 10.3390/cancers13112783

17 Remšík, J. et al. Plasticity and Intratumoural Heterogeneity of Cell Surface Antigen Expression in Breast Cancer. British Journal of Cancer 118, 813–819 (2018). 10.1038/bjc.2017.497

18 Dutta, S. et al. Angiogenin interacts with the plasminogen activation system at the cell surface of breast cancer cells to regulate plasmin formation and cell migration. Mol Oncol 8, 483–507 (2014). 10.1016/j.molonc.2013.12.017

19 Liu, G. et al. Glutamate dehydrogenase is a novel prognostic marker and predicts metastases in colorectal cancer patients. J Transl Med 13, 144 (2015). 10.1186/s12967-015-0500-6

20 Noh, H., Hong, S. & Huang, S. Role of urokinase receptor in tumor progression and development. Theranostics 3, 487–495 (2013). 10.7150/thno.4218

21 Liu, J.-F., Chen, P.-C., Chang, T.-M. & Hou, C.-H. Monocyte Chemoattractant Protein-1 promotes cancer cell migration via c-Raf/MAPK/AP-1 pathway and MMP-9 production in osteosarcoma. Journal of Experimental & Clinical Cancer Research 39, 254 (2020). 10.1186/s13046-020-01756-y

22 Wolczyk, D. et al. TNF-α promotes breast cancer cell migration and enhances the concentration of membrane-associated proteases in lipid rafts. Cell Oncol (Dordr) 39, 353–363 (2016). 10.1007/s13402-016-0280-x

23 Mercogliano, M. F., Bruni, S., Elizalde, P. V. & Schillaci, R. Tumor Necrosis Factor α Blockade: An Opportunity to Tackle Breast Cancer. Front Oncol 10, 584 (2020). 10.3389/fonc.2020.00584

24 Sheikhpour, E. et al. A Survey on the Role of Interleukin-10 in Breast Cancer: A Narrative. Rep Biochem Mol Biol 7, 30–37 (2018).

25 Eyre, R. et al. Microenvironmental IL1β promotes breast cancer metastatic colonisation in the bone via activation of Wnt signalling. Nature communications 10, 5016 (2019). 10.1038/s41467-019-12807-0

26 Choi, J. et al. Inflammatory Signals Induce AT2 Cell-Derived Damage-Associated Transient Progenitors that Mediate Alveolar Regeneration. Cell Stem Cell 27, 366–382.e367 (2020). 10.1016/j.stem.2020.06.020

27 Liang, Y. et al. CX3CL1 involves in breast cancer metastasizing to the spine via the Src/FAK signaling pathway. J Cancer 9, 3603–3612 (2018). 10.7150/jca.26497

28 Shang, G. S., Liu, L. & Qin, Y. W. IL-6 and TNF-α promote metastasis of lung cancer by inducing epithelial-mesenchymal transition. Oncol Lett 13, 4657–4660 (2017). 10.3892/ol.2017.6048

29 Reeves, M. E. et al. Ras-Association Domain Family 1C Protein Promotes Breast Cancer Cell Migration and Attenuates Apoptosis. BMC cancer 10 (2010). 10.1186/1471-2407-10-562

30 Pein, M. et al. Metastasis-initiating cells induce and exploit a fibroblast niche to fuel malignant colonization of the lungs. Nature communications 11, 1494 (2020). 10.1038/s41467-020-15188-x

31 Bach, D.-H., Park, H. J. & Lee, S. K. The Dual Role of Bone Morphogenetic Proteins in Cancer. Molecular Therapy - Oncolytics 8, 1–13 (2018). 10.1016/j.omto.2017.10.002

32 Doyle, L. M. & Wang, M. Z. Overview of Extracellular Vesicles, Their Origin, Composition, Purpose, and Methods for Exosome Isolation and Analysis. Cells 8 (2019). 10.3390/cells8070727

33 Benedikter, B. J. et al. Ultrafiltration combined with size exclusion chromatography efficiently isolates extracellular vesicles from cell culture media for compositional and functional studies. Scientific reports 7, 15297 (2017). 10.1038/s41598-017-15717-7

34 Ondruššek, R. et al. Prognostic value and multifaceted roles of tetraspanin CD9 in cancer. Front Oncol 13, 1140738 (2023). 10.3389/fonc.2023.1140738

35 Tang, Q. et al. Recent Advances in Detection for Breast-Cancer-Derived Exosomes. Molecules 27 (2022). 10.3390/molecules27196673

36 Ramos, E. K. et al. CD81 partners with CD44 in promoting exosome biogenesis, tumor cluster formation, and lung metastasis in triple negative breast cancer. bioRxiv, 2022.2002.2023.481674 (2022). 10.1101/2022.02.23.481674

