## Supplementary Figures and Table for "An *in vitro* model of breast cancer metastatic niche priming"

**Supplemental Figures and Tables.**

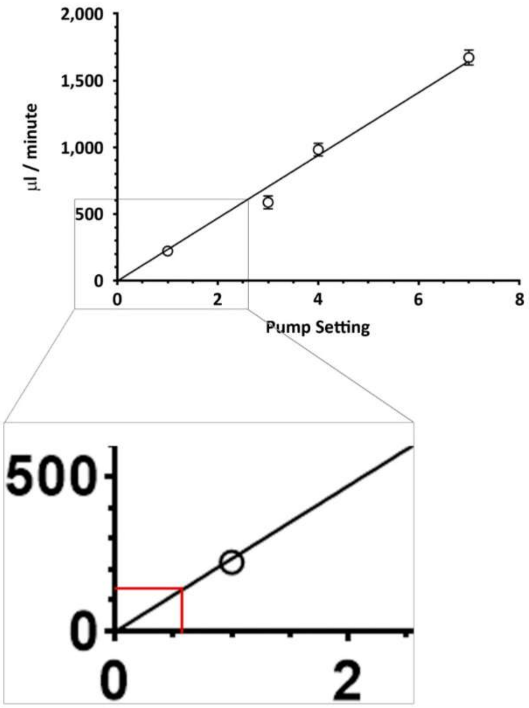


**Supplemental Figure 1: Linked culture can mimic tail vein injection used in the Pulmonary Metastasis Assay.**

The *Quasi vivo* circuit must be calibrated before use to identify the correct setting to achieve the required µL / minute rate. The required number of chambers are linked, and the circuit is filled with PBS. Rather than returning the liquid to the reservoir, the PBS is allowed to flow into a weigh boat. The circuit is run, with various pump settings, for 2 minutes and the PBS expelled is weighed on a fine balance. This measurement is performed 3 times for each pump setting.

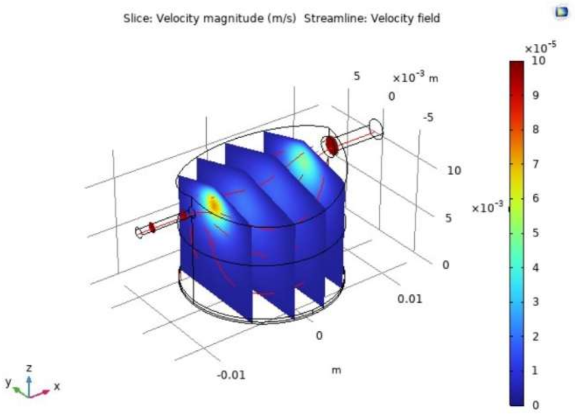


**Supplemental Figure 2: Shear stress within chambers.**

The shear stress on the base is very low, with an average of 0.486 microPa and a maximum 1.34 microPa.

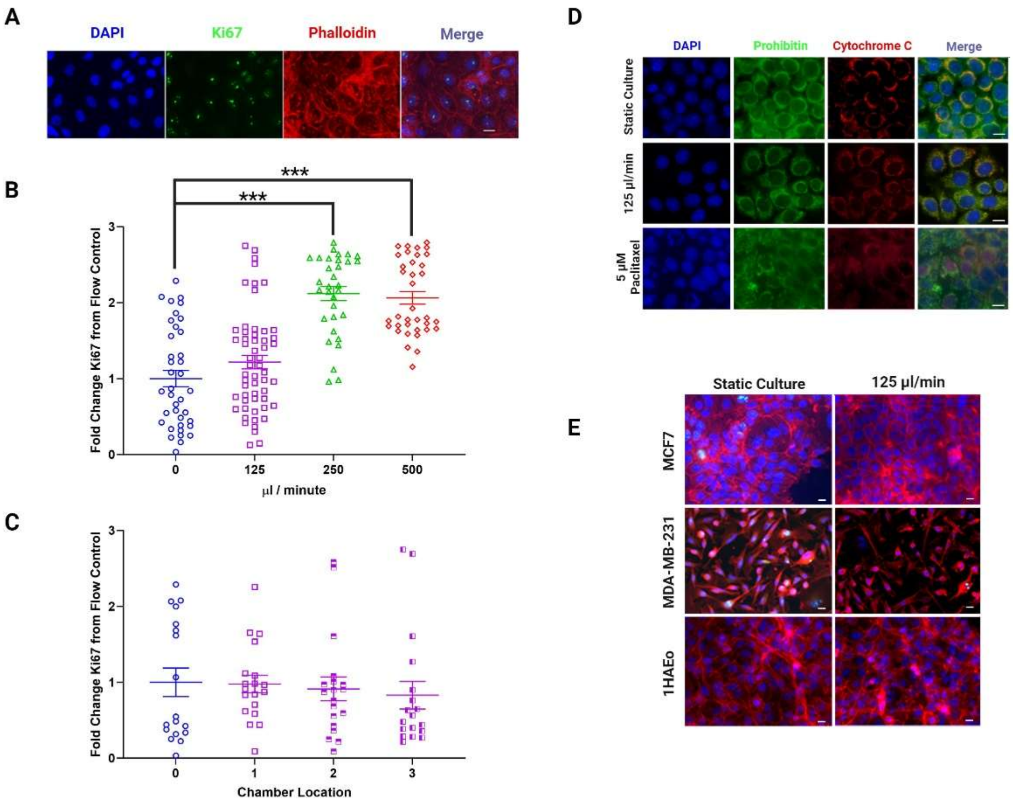


**Supplemental Figure 3: Flow rate.**

A) Ki67 was assessed using immunofluorescence counting positive cells as a percentage of all cells in 5 fields of view. B) Flow rates of 250 μL/minute and above resulted in increased proliferation. C) No difference was seen between multiple chambers. D) Representative images of cells stained with Cytochrome C to assess release into the nucleus in apoptotic cells, 5 μM Paclitaxel was used as a control. E) No change in morphology was detected using Phalloidin cell membrane stain. Scale bar in each image represents 10μm.

One-way ANOVA with Dunnett's correction, ***$P$<0.001

| **These items are the basic minimum to include in a manuscript. Without this information, readers and reviewers cannot assess the reliability of the findings.** | | | |
|---|---|---|---|
| **Item** | | **Recommendation** | **Section/line number, or reason for not reporting** |
| **Study design** | 1 | For each experiment, provide brief details of study design including:<br>**a)** The groups being compared, including control groups. If no control group has been used, the rationale should be stated.<br>**b)** The experimental unit (e.g. a single animal, litter, or cage of animals). | **a)** Mean fluorescence was assessed at day 7 and compared to day 0. As there was no "test" performed no control group was required.<br><br>**b)** Experimental unit was lung slice. **Page 13.** |
| **Sample size** | 2 | **a)** Specify the exact number of experimental units allocated to each group, and the total number in each experiment. Also indicate the total number of animals used.<br>**b)** Explain how the sample size was decided. Provide details of any *a priori* sample size calculation, if done. | **a)** 2.9.2 Ex vivo lung samples, **page 12** – 6 mice TOTAL, including:<br>2.9.2.1 Pulmonary metastasis assay, **Page 12-13** – 3 mice<br>2.9.2.2 Homing and colonisation to PuMA slices., **page 13** – 3 mice<br><br>**b)** Calculations were made based upon previous work and suggested 12-15 samples (lung slices) were needed for each group to achieve 80% power with $P<0.05$. |
| **Inclusion and exclusion criteria** | 3 | **a)** Describe any criteria used for including and excluding animals (or experimental units) during the experiment, and data points during the analysis. Specify if these criteria were established *a priori.* If no criteria were set, state this explicitly.<br>**b)** For each experimental group, report any animals, experimental units or data points not included in the analysis and explain why. If there were no exclusions, state so.<br>**c)** For each analysis, report the exact value of *n* in each experimental group. | **a)** Slices which were not complete/too thick were excluded. Lungs which were not successfully perfused were not used.<br><br>**b)** All data points are included<br><br>**c)** n = 5 slices from 3 mice in each case. Figure 7 legend, **page 26** |
| **Randomisation** | 4 | **a)** State whether randomisation was used to allocate experimental units to control and treatment groups. If done, provide the method used to generate the randomisation sequence.<br>**b)** Describe the strategy used to minimise potential confounders such as the order of treatments and measurements, or animal/cage location. If confounders were not controlled, state this explicitly. | **a)** Lung slices we prepared from all animals and randomly assigned to day 0 or day 7 – **Page 13**<br><br>**b)** Scoring was performed by AI meaning blinding was not necessary but scanned images were deidentified so that training of AI was not performed on a specific sub-set. |
| **Blinding** | 5 | Describe who was aware of the group allocation at the different stages of the experiment (during the allocation, the conduct of the experiment, the outcome assessment, and the data analysis). | Slides were scanned by staff in imaging core facility and, although slide labels were included in the image, the details were hidden from view during analysis. |

| | | | |
|---|---|---|---|
| **Outcome measures** | **6** | **a) Clearly define all outcome measures assessed (e.g. cell death, molecular markers, or behavioural changes).**<br>**b) For hypothesis-testing studies, specify the primary outcome measure, i.e. the outcome measure that was used to determine the sample size.** | **a)** Mean fluorescence area was assessed for each scanned lung using confocal microscopy<br><br>**b)** N/A |
| **Statistical methods** | 7 | **a)** Provide details of the statistical methods used for each analysis, including software used.<br>**b)** Describe any methods used to assess whether the data met the assumptions of the statistical approach, and what was done if the assumptions were not met. | **a)** GraphPad Prism was used for analysis and unpaired t-tests were performed on raw data. Details in 2.10 Statistics, **page 13** and in Figure 7 legend, **page 26.**<br><br>**b)** Normality and descriptive statistics were used to confirm distribution and variance. T-test is selected based on these data. |
| **Experimental animals** | 8 | **a)** Provide species-appropriate details of the animals used, including species, strain and substrain, sex, age or developmental stage, and, if relevant, weight.<br>**b)** Provide further relevant information on the provenance of animals, health/immune status, genetic modification status, genotype, and any previous procedures. | Details are provided in 2.9.2 Ex vivo lung samples, **page 12.**<br>- NOD SCID gamma mice<br>- Charles River<br>- female (relating to breast cancer which is predominantly a female disease)<br>- 8-10 weeks<br>- over 18 grams |
| **Experimental procedures** | 9 | For each experimental group, including controls, describe the procedures in enough detail to allow others to replicate them, including:<br>• What was done, how it was done and what was used.<br>• When and how often.<br>• Where (including detail of any acclimatisation periods).<br>• Why (provide rationale for procedures). | Full details are provided in 2.9.2.1 Pulmonary metastasis assay. **page 12.** |
| **Results** | 10 | **a)** For each experiment conducted, including independent replications, report: summary/descriptive statistics for each experimental group, with a measure of variability where applicable (e.g. mean and SD, or median and range).<br>**b)** If applicable, the effect size with a confidence interval. | See figure 7<br>a) Data was converted to Log2 fold change so that increase and decrease were considered equally and so that comparisons could be made between the higher starting cell number following tail vein injection and the lower levels following linked culture.<br><br>**b)** N/A |
| **These items complement the Essential 10 set and add important context to the study described. Reporting the items in both sets represents best practice.** | | | |
| **Abstract** | **11** | **Provide an accurate summary of the research objectives, animal species, strain and sex, key methods, principal findings, and study conclusions.** | Word limit and the minimal use of animals means we do not reference mouse details in the abstract. |

| | | | |
|---|---|---|---|
| **Background** | **12** | **a) Include sufficient scientific background to understand the rationale and context for the study and explain the experimental approach.**<br>**b) Explain how the animal species and model used address the scientific objectives and, where appropriate, the relevance to human biology.** | **a)** Discussion of the limitations of in vivo modelling are included in the introduction as this forms the argument for need of the model we have developed. **Pages 3-5**<br>b) PuMA model is described and discussed in methods. 2.9.2 Ex vivo lung samples. **Pages 12-13** |
| **Objectives** | 13 | Clearly describe the research question, research objectives and, where appropriate, specific hypotheses being tested. | See 1. Introduction and 2.9.2.1 Pulmonary metastasis assay. |
| **Ethical statement** | 14 | Provide the name of the ethical review committee or equivalent that has approved the use of animals in this study and any relevant license or protocol numbers (if applicable). If ethical approval was not sought or granted, provide a justification. | See 2.9.2 Ex vivo lung samples. |
| **Housing and husbandry** | 15 | Provide details of housing and husbandry conditions, including any environmental enrichment. | Cages - Emerald line IVC EM500<br>Housing - Animals are group housed<br>Water - Automatic RO water<br>Bedding - Datesand Aspen Chip 2<br>Enrichment - Sizzlenest, Happimat, Wooden chew stick, Polycarbonate mouse tent, Polycarbonate handling tube, Loft<br>Light cycle - 12:12 6am -6pm<br>Temp - 20-24°C<br>Humidity - 45-65% RH |
| **Animal care and monitoring** | 16 | **a)** Describe any interventions or steps taken in the experimental protocols to reduce pain, suffering, and distress.<br>**b)** Report any expected or unexpected adverse events.<br>**c)** Describe the humane endpoints established for the study, the signs that were monitored, and the frequency of monitoring. If the study did not have humane endpoints, state this. | **a)** Using appropriate needle size: 27G or smaller, new needle per mouse, warming the mouse prior to injection to dilate the blood vessel. Used overdose of Isoflurane to cull the mouse and cessation of circulation (cutting the femoral) to confirm death. It is the most appropriate method of culling to keep the trachea intact to allow for the lungs to be filled.<br>**b)** N/A, none<br>**c)** N/A, endpoints occurred *in vitro*. |
| **Interpretation/ scientific implications** | 17 | **a)** Interpret the results, taking into account the study objectives and hypotheses, current theory, and other relevant studies in the literature.<br>**b)** Comment on the study limitations including potential sources of bias, limitations of the animal model, and imprecision associated with the results. | See results and discussion |
| **Generalisability/ translation** | **18** | **Comment on whether, and how, the findings of this study are likely to generalise to other species or experimental conditions, including any relevance to human biology (where appropriate).** | **Pages 26 - 29** – discussion of replacement opportunity |

| | | | |
|---|---|---|---|
| **Protocol registration** | **19** | **Provide a statement indicating whether a protocol (including the research question, key design features, and analysis plan) was prepared before the study, and if and where this protocol was registered.** | Protocol was not registered. |
| **Data access** | 20 | Provide a statement describing if and where study data are available. | No data was produced that is not displayed within the body of the paper**,** so availability is not relevant. |
| **Declarations of Interests** | **21** | **a)** Declare any potential conflicts of interest, including financial and non-financial. If none exist, this should be stated.<br>**b)** List all funding sources (including grant identifier) and the role of the funder(s) in the design, analysis, and reporting of the study. | **a)** See Title page, **page 1**<br>**b)** See Acknowledgements**, page 30** |

**Supplemental Table 1: Arrive guidelines were adhered to during this project**
